# DUALISTIC SAHA DOSE-DEPENDENT EFFECTS ON GLIAL-INFLAMMATORY RESPONSE

**DOI:** 10.1101/2024.01.16.575689

**Authors:** S. Mancino, M. Boraso, A. Galmozzi, E. De Fabiani, Crestani, B. Viviani

## Abstract

Neuroinflammation comprises biochemical and cellular responses of the nervous system to injury, emerging as a central process in many neurological conditions. Although beneficial in nature, over-activation of the immune response may result in the production of neurotoxic factors that exacerbate the disease state. Due to the relevant role of histone acetylation in transcriptional regulation, inhibition of histone deacetylase (HDAC) activity plays a central role in the pathogenesis of inflammation. Here, we investigated the impact of HDACs inhibition in the regulation of the inflammatory response elicited by lipopolysaccharide (LPS) in glia cells using trichostatin A (TSA) and suberoylanilide hydroxamic acid (SAHA). We observed that acute and prolonged exposure to low doses of TSA boosts the inflammatory response in glia cells. Conversely, TSA pre-treatment in cells exposed to LPS decreases inflammation. Additionally, we observed that low and high doses of SAHA exert opposite effects on LPS-induced inflammatory response. Low dose (100 nM) potentiates inflammatory response by increasing the production of pro-inflammatory cytokines tumour necrosis factor (TNF)-α and interleukin (IL) - 1β; and reducing the anti-inflammatory mediator IL-10. Nevertheless, 5 µM SAHA reverts the pro-towards anti-inflammatory profile.

To better characterize SAHA dualistic effects in glia cells, we generated comparative genome-wide gene expression data. Simultaneous exposure to 5 µM SAHA and LPS (10 ng/ml) alters the expression of 1628 genes, while the combination of 100 nM SAHA and LPS regulates the expression of 97 transcripts. Both doses of SAHA differentially regulate important pathways involved in cellular function maintenance, homeostasis, and survival.

Likewise, inhibition of the Janus Kinase - signal transducers and activators of the transcription (JAK/STAT) signalling pathway seems to mediate the anti-inflammatory effects of 5 µM SAHA. On the other hand, 100 nM SAHA increases pro-inflammatory cytokine levels possibly via modulation of the underlined feedback mechanisms triggered by IL-10 expression regulation.

Together, these results contribute to outline a comprehensive picture of the involvement of HDACs inhibitors (HDACi) in the onset or prevention of neuroinflammation, showing that their effects depend on cell types, HDACi dosage and specificity, and protocol used.

## 1. INTRODUCTION

Marked inflammatory reaction in the central nervous system (CNS) is involved in the pathogenesis, progression, and exacerbation of neurodegenerative diseases [1] and psychiatric disorders [2]. Epigenome modifications, such as histone modifications [3], lie behind the induction and maintenance of the inflammatory state. Indeed, acetylation and deacetylation of lysine residues is involved in the regulation of gene expression through modification of chromatin conformation, ultimately affecting several cellular processes including inflammation [4]. These post-translational modifications are regulated by the activity of two classes of enzymes: histone acetyltransferases (HATs) and histone deacetylase (HDACs), respectively. HATs, by adding an acetyl group to lysine of core histone, lead to a relaxed chromatin structure generally correlated with gene activation. HDACs, instead, revert the lysine acetylation to stabilize chromatin architecture and usually mediate transcriptional repression [5]. The levels and activities of HATs and HDACs are finely balanced in neuronal cells under normal conditions [6]. However, in neurodegenerative state, the histone acetylation homeostasis is greatly impaired, shifting towards hypoacetylation [6]. Thus, the inhibition of HDACs has been attempted to restore homeostatic conditions.

Eighteen members of HDAC family divided in four classes have been identified in humans. class I (HDACs 1, 2, 3, and 8), class IIA (HDACs 4, 5, 7, and 9), class IIB (HDACs 6 and 10), class III (Sirtuins 1 to 7), and class IV (HDAC 11) [7]. HDAC 2, 4, 5, and 6 play a key role in the regulation of genes involved in inflammation [8]. Class I HDACs are mainly involved in innate immunity by regulation of inflammatory reactions while class II HDACs are associated with adaptive immunity [10]. As such, the potential use of HDACs in inflammatory diseases [9] is currently under discussion for their ability to target many immune cell types [10] [11], including macrophages [12].

Suberoylanilide hydroxamic acid (SAHA) and Trichostatin A (TSA) are both nonselective small molecule inhibitors of the Zn (II)-dependent class I and class II HDACs, considered suitable therapeutics for various malignant diseases and few HDACi are also considered for the treatment of neurodegenerative diseases [13]. Indeed, SAHA, also known as Vorinostat, is approved in some countries for the treatment of cutaneous T-cell lymphoma [14], it is in clinical testing for other cancer types [15], HIV [16] and is under evaluation to determine the maximal tolerable dose in the treatment of Alzheimer’s Disease (NCT03056495).

Despite this evidence, very little is known about the regulation of inflammatory processes driven by HDACs in the CNS and how it affects the homeostasis and function of immune resident cells such as microglia [17] [18] and astrocytes [19]. The histone deacetylases HDAC 1 and HDAC 2 are considered critical regulators of microglia development, homeostasis, maturation, and activation [20]. However, their involvement in the molecular mechanisms underlying neuroinflammation is not fully elucidated and conflicting results are reported [21][22]. Indeed, *in vitro* studies suggest a different regulation of the inflammatory process by histone deacetylase inhibitors (HDACi). This may depend on cell types, reporting a potentiation of the inflammatory response by HDACi in mouse microglia cell lines [23], primary rat microglia [20][24] and macrophages [25], while a decrease of inflammation in primary mouse dendritic cells [26], mixed CNS glia [27], primary mouse microglia, hippocampal HT-22 cells [28], and macrophages [29][11].

Moreover, contradictory results in the regulation of neuroinflammation by HDACi were obtained, contingent to the experimental protocol used *in vitro* studies. Pre-treatment with TSA or SAHA before the inflammatory stimulus, reduces the production of pro-inflammatory cytokines in primary culture of microglia cells [30][31]. On the other hand, TSA and SAHA treatments enhance the inflammatory response of LPS when added simultaneously to the inflammatory trigger in primary cultures of microglia cells [23][20][24]. All together this evidence shows a dualistic effect of HDACs inhibitors on inflammatory response that could be ascribed to different factors, such as the dose, protocol and cell type used. To clarify the modulatory effects of HDACi under different conditions, the first part of this study aimed to investigate the ability of HDACi to modulate inflammation by comparing the response between different cellular populations involved in central and peripheral inflammatory response (i.e. mixed glial cells, purified microglia and astrocytes, alveolar macrophages), using different paradigms of treatment (simultaneous or prior treatment with pro-inflammatory stimulus) and different HDACi, wide (TSA or SAHA) or class I (MS275) and class II (MC1568) selective spectrum.

The second part of this study, using a global transcriptomic approach, uncovered gene networks and pathways that are affected by experimental treatment conditions leading to an opposite modulation of LPS-induced response (100 nM or 5 µM SAHA) in glial cells. Our findings provide a comprehensive picture of the gene pathways and signalling events altered using HDACi during the inflammatory response and support a potential role of SAHA in the onset or prevention of neuroinflammation.

## 2. MATERIALS AND METHODS

### 2.1 Reagents

All cell reagents, unless otherwise specified, were purchased from Sigma-Aldrich Co. (St. Louis, MO, USA) at the highest purity available. Lipopolysaccharide (LPS) from *Escherichia coli* serotype 0127: B8 was obtained from Sigma-Aldrich Co. and the stock solution was dissolved in PBS to a final concentration of 5 mg/ml. Tricostatin A was purchased from Sigma-Aldrich Co., Suberoylanilide hydroxamic acid was purchased from Cayman (Michigan, USA), MS275 and MC1568 were synthetized and were kind gifts by Antonello Mai (“Sapienza” Università di Roma). The stock solutions of inhibitors were dissolved in DMSO to a final concentration of 20 mM (final concentration of DMSO in culture medium <0.1%). Reagents and primers (18s: Hs99999901_s1; TNF-α: Rn99999017_m1; IL-1 β: Rn00580432_m1; IL-10: Rn00563409_m1) for real time reverse transcriptase polymerase chain reaction were from Applied Biosystems (Foster City, CA, USA).

### 2.2 Cell cultures

#### Primary culture of glia cells

Primary cultures of glial cells were prepared from 2-day-old newborn rats (Sprague–Dawley, Charles River, Calco, Italy), following a standardized procedure [32][33]. Experimental procedures were conducted following the National Institutes of Health Guide for the Care and Use of Laboratory Animals, the European Community Council Directives 2010/63/EU, and the Italian law 26/2014. Approval was obtained from the local Animal Use Committee and the Italian Ministry of Health (permits 475/2015 PR).

Cerebral hemispheres were freed of the meninges and were mechanically disrupted; cells were dispersed in a solution of trypsin 2.5% and 1 mg/ml DNAse, filtered through a 100-um nylon mesh. Cells were seeded in 24-well plates for the treatments of mixed glia cells (5×10^4^ cells per well) and in a 75 cm^2^ flask for the preparation of microglia and astrocytes (5×10^6^ cells per flask), in Minimum essential Eagle’s Medium (MEM) supplemented with 10% fetal calf serum (FCS), 0.6%glucose, 0.1 mg/ml streptomycin, 100 IU/ml penicillin and 2 mM L-glutammine. Glial cultures were fed twice a week and grown up to confluence at 37 °C in a humidified incubator with 5% CO_2_. Cells were treated after 10-15 days from plating when they reached confluence.

#### Primary cultures of astrocytes and microglia cells

A layer of astrocytic cells was obtained through vigorous shaking of a confluent 10-day-old monolayer of mixed glial cells, as described by [34]. Cultures of enriched astroglia were treated further with 5 mM L-leucine methyl ester to eliminate microglia (97% homogeneity). Isolated astroglial preparations were then plated in 24-well plates (10^5^ cells per well) in MEM with supplements as described above and treated two days after seeding. Microglia were isolated by shaking glial cultures at 800 x *g* for 2 hours. Microglia, which dislodged into the medium, were purified by plating for 30 min in 24-well plates (10^5^ cells per well) in MEM with supplements as above and FCS 15%. Contaminating cells were removed with supernatant. These conditions allowed us to obtain highly enriched microglial cultures with 98% homogeneity, as assessed by immunocytochemistry with *Griffonia simplicifolia* isolectin B_4_. The treatments of microglia were performed the day after the plating.

#### Primary culture of alveolar macrophages

Alveolar macrophages were collected by bronchoalveolar lavage as previously described [35]. Recovery was 10-15×10^6^ cells per rat, of which > 98% were macrophages, as assessed by Giemsa stain. Once washed and resuspended to 10^6^ viable alveolar macrophages/ml, the cells were seeded in 24-well plates (3×10^5^ cells per well) and were allowed to adhere to plastic plates in RPMI 1640 containing 10% FCS, streptomycin (0.1 mg/ml), penicillin (100 International Units/ml), L-glutammine (2 mM) and gentamicin (50 ng/ml) for 1 hour at 37° in 5% CO_2_, and then treated.

### 2.3 Viability Assay

Cell viability was measured by the 3-(4,5-dimethyl-thiazol-2-yl)-2,5-diphenyltetrazolium bromide (MTT) assay [36]. MTT tetrazolium salt was dissolved in serum-free medium to a final concentration of 0.75 mg/ml and added to the cells (glia or macrophages) after 24 hours of treatment with ± 10 ng/ml LPS ± 100 nM TSA. Cells were then incubated for 3 hours at 37 °C. The medium was then removed and formazan extracted with 1N HCl:isopropyl alcohol (1:24). Absorbance of the resulting solutions was read at 595 nm in a microplate reader (Molecular Devices, Sunnyvale, CA). Results are expressed as the percentage of control cells ± Standard Error of the Means (SEM).

### 2.4 RNA extraction and real time Reverse Transcriptase-Polymerase Chain Reaction (RT-PCR)

Total RNA was isolated from glia cells after 1, 3 and 6 hours of treatment with 10 ng/ml LPS and with or without 10 nM TSA, using a commercially available kit (TriReagent from Sigma) as described by the manufacturer. For the synthesis of cDNA, 2 μg of total RNA was reverse-transcribed using a high-capacity cDNA archive kit from Applied Biosystems (Foster City, CA, USA) following the supplier’s instructions. TNF-α, IL-1β and IL-10 gene expression were evaluated by real time reverse transcription polymerase chain reaction (Real Time RT-PCR). For PCR-analysis, Taq-Man™-PCR technology was used. For each PCR reaction, 40 ng of total RNA were used. The 18S ribosomal RNA was used as an endogenous reference and the quantification of the transcripts was performed by the ΔΔC_T_ method.

### 2.5 Biological assay for TNF-α

TNF-α content in the medium obtained from treated cells was assayed by determining the cytotoxicity of TNF-α against sensitive L929 cells (mouse fibroblast) [35]. Briefly, L929 cells were seeded into 96-well microtitre plates (2.5×10^4^ cells per well) in RPMI-1460 culture medium supplemented with 10% FCS, streptomycin (0.1 mg/ml), penicillin (100 International Units/ml), L-glutammine (2 mM) and incubated for 24 hours at 37 °C in a humidified atmosphere of 5% CO_2_. After removal of the medium, actinomycin D (0.6 μg/μl) was added to each well for 1 hour at 37 °C. Recombinant murine TNF-α and test samples were added, and plates were incubated at 37°C for 18 hours. Cells were stained and fixed with 0.2% crystal violet in 2% ethanol for 10 min at room temperature and then lysed with 1% sodium dodecyl sulfate. Absorbance at 505 nm was detected with a microplate reader (Molecular Devices, Sunnyvale, CA). TNF-α concentration was calculated from a standard curve based on known amounts of recombinant murine TNF-α. Data were normalized to LPS and presented as percentage change in TNF-α levels ± SEM.

### 2.6 IL-1β and IL-10 assay

IL-1β and IL-10 amounts were measured in the cultured media of treated cells by commercially available ELISA kit as described by manufacture (Quantikine, R&D Systems, Abingdon, UK). Data were normalized to LPS and presented as percentage change in cytokines release ± SEM.

### 2.7 Microarray data analysis

Total RNA was isolated from the glia cells using the RNeasy Kit (Qiagen, Milano, Italy) following the manufacturer’s instructions for cells treated with 100 nM or 5μM SAHA in the absence or presence of 10 ng/ml LPS after 4 hours.

RNA quality was established using Bioanalyzer (Agilent). Synthesis of cDNA was carried out starting from 5 µg of total RNA.

Microarray analysis was performed at Genopolis Consortium of Functional Genomic, Università di Milano – Bicocca, using a standard protocol described in [37].

#### Sample treatments

Different cell conditions were examined: glia cells without treatment/induction (control, CTRL), glia cells induced with 10 ng/ml LPS (LPS), glia cells treated with 100 nM SAHA or 5 μM SAHA (SAHA100 or SAHA5), glia cells treated with 100 nM SAHA or 5 μM SAHA with a simultaneous LPS induction (LPS_SAHA100 or LPS_SAHA5).

For each condition, three samples were considered for the analysis of gene expression by AMDA microarray technology [37].

#### Microarray comparisons and DEG analysis

Differentially expressed genes (DEGs) were identified comparing gene expression profiles of glia cells treated with 100 nM SAHA or 5 μM SAHA in the presence or the absence of the inflammatory stimulus. Comparisons are reported in:

- Table S1 (Supplementary Materials), LPS_SAHA100 vs LPS; LPS_SAHA5 vs LPS,
- Table S2 (Supplementary Materials), SAHA100 vs CTRL; SAHA5 vs CTRL,
- Venn diagrams (Fig S4), SAHA100_CTRL vs SAHA100_LPS, SAHA5_CTRL vs SAHA5_LPS).

This multivariate analysis aims to reveal relationships between genes in the different treatment paradigms considered.

The identification of DEGs was addressed using Linear Models for Microarray Data (LIMMA). For the detection, linear modeling approach and empirical Bayes methods together with false discovery rate correction of the p-value (Benjamini-Hochberg) were used to moderate the standard errors. All the requested comparisons were performed selecting DEG with a threshold p Value of 0.01 as the criterion for statistical significance with a fold-change threshold of 1.

#### 2.7.1 Analysis of data and canonical pathway enrichment analyses

DEGs significantly (log 2-Fold Change and P values) up- or downregulated genes were associated with canonical pathways. A functional annotation of DEG is performed based on a subset of the annotation provided by the Bioconductor project (www.bioconductor.org), [37].

##### Functional annotation

Canonical pathway enrichment analyses were performed using Kyoto encyclopedia of genes and genome (KEGG) pathways as well as Gene Ontology (GO) enrichment analyses (www.genome.jp/kegg, www.geneontology.org respectively). The most representative functional annotations for DEGs from each experimental condition were identified and only the top 40 statistically most significant annotation terms are reported. If p-values were lower than 10^-5^, the enrichment was considered significant [37].

##### Enrichment analysis

Only pathways represented with at least 2 DEGs were ranked according to their relevance. Based on this ranking only the first 40 (or less whenever this number was not reached) were reported. If at least two experimental conditions were analyzed, a functional summary reporting the enrichment p-values less than 0.05 (log10 of p-value) for the different annotation terms and their similarity is provided [37].

Tables embedded in the Supplementary Information contain lists of the relevant annotation terms for each cluster and for each annotation source and the functional p value enrichment.

### 2.8 Statistical analysis

All experiments were performed on at least two different primary cell preparations. Data are expressed as mean ± SEM. Data were analyzed using GraphPad Prism 9 edition (GraphPad Software, La Jolla, CA, USA). Statistical differences were determined using analysis of variance followed by multiple comparison test, as indicated in the legends. Effects were designated significant when *p*<0.05.

## 3. RESULTS

### 3.1 Dose-dependent effects of TSA on the expression of pro- and anti-inflammatory cytokines in LPS-co-stimulated mixed and purified glia cells

To address how broad-spectrum inhibition of HDACs modulates the neuroinflammatory response in mixed glia *in vitro*, primary cultures of glia cells were simultaneously exposed to bacterial LPS (10 ng/ml) and increasing concentrations of TSA (0.01, 0.1, 1, 10, and 100 nM, respectively) for 24 hours (Fig 1 A). The release of TNF-α and IL-1β (early phase pro-inflammatory cytokines) and IL-10 (late phase anti-inflammatory cytokine) was evaluated in the culture medium by a specific biological assay or ELISA, respectively (Fig 1 B). The selected concentration of LPS of 10 ng/ml allowed to obtain the highest cellular response without affecting their viability after 24 hours of treatment (cell viability: control = 100% ± 10.4, LPS 10 ng/ml = 89.9% ± 10.3). Moreover, all the tested concentrations of TSA, with or without LPS 10 ng/ml, did not affect glial cell viability (cell viability referred to TSA highest concentration: control=100% ± 3.6, TSA 100 nM = 116.7 ± 4.6, LPS 10 ng/ml + TSA 100 nM = 86.7% ± 9.2). As reported in Fig 1 B, TSA enhanced LPS-induced release of the pro-inflammatory cytokines TNFα (Fig 1 B, inset i) and IL–1β (Fig 1 B, inset ii). The effect was dose-dependent and maximal at the concentration of 10 nM TSA while became less marked at TSA of 100 nM, (Fig 1 B, inset i and ii). On the contrary, TSA reduced the production of IL-10 elicited by LPS treatment up to 10 nM (Fig 1B, inset iii). However, at the concentration of 100 nM, TSA significantly reverted the trend and increased the production of the anti-inflammatory mediator IL-10 (Fig 1 B, inset iii). Treatment with TSA itself did not induce the release of TNF-α, IL-1β, or IL-10 (data not shown).

**Figure 1.**
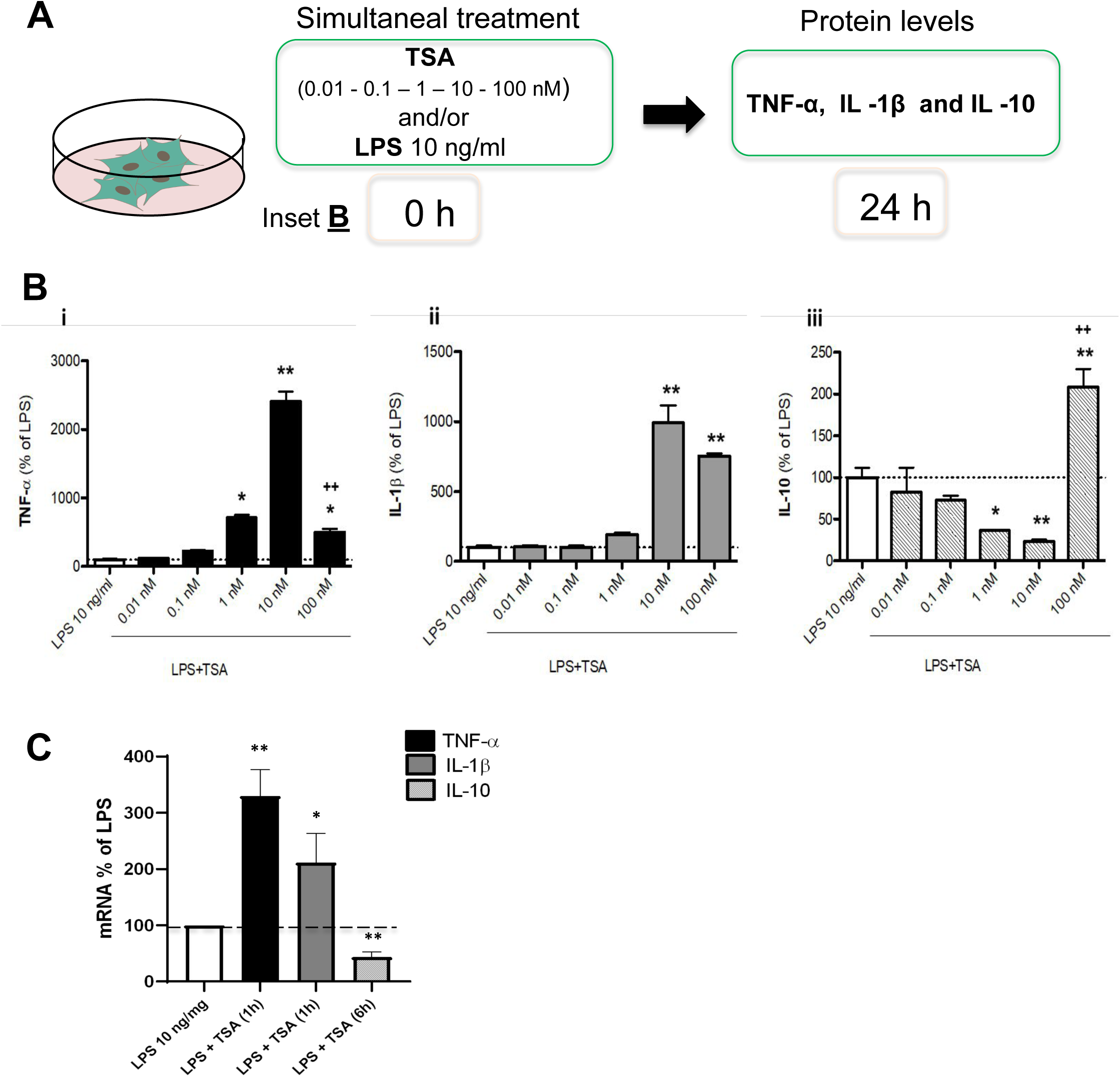
Effect of exposure to TSA on cytokines release (B) and mRNA cytokines expression (C) in simultaneous LPS-stimulated glial cells. **A)** Experimental design for prolonged (24 hours) TSA exposure. **B)** Prolonged (24 hours) glia cells exposure to 10 ng/ml LPS in absence or presence of different TSA concentrations (0.01, 0.1, 1, 10, 100 nM) to evaluate: TNF-α **(i**), IL - 1β **(ii)** and IL - 10 **(iii)** concentrations, for the condition described above, measured in the culture medium by biological assay or ELISA. The results are expressed as a percentage of LPS. **C)** Short (1 and 6 hours) glia cells exposure to 10 ng/ml LPS in absence or simultaneous presence of 10 nM TSA to evaluate mRNA levels of TNF-α (black bars), IL - 1β (grey bars) and IL - 10 (stripped bars) by RT-PCR. The results are expressed as a percentage of LPS. Data are mean ± SEM of two or three independent experiments in triplicate. Statistical analysis was performed with one-way ANOVA followed by Tukey’s test, \**p*<0.05, \*\**p*<0.01 LPS+TSA *vs* LPS, ^++^ *p*<0.01 LPS+TSA 10 nM *vs* LPS+TSA 100 nM, and LPS+TSA 10 nM *vs* LPS.

Mixed glia cells were also treated with 10 ng/ml LPS in the absence or presence of 10 nM TSA for 1 hour (early phase pro-inflammatory) and 6 hours (late phase anti-inflammatory) and the level of cytokines mRNA were evaluated by RT-PCR (Fig 1 C). Accordingly, after 1h, 10 nM TSA enhanced LPS-induced mRNA expression of pro-inflammatory cytokines TNF-α and IL-1 β, and significantly reduced LPS-induced mRNA of the anti-inflammatory cytokine IL-10 at 6h. (Fig 1 C).

Purified primary cultures of microglia and purified astrocytes were treated with 10 ng/ml LPS with or without 10 nM TSA for 24 hours. At the end of the treatment, the release of TNF-α, IL-1β, and IL-10 in the culture medium was evaluated. Consistent with the data obtained in the mixed glia population, 10 nM TSA significantly potentiated LPS-induced release of the two pro-inflammatory cytokines TNF-α and IL-1β in both glia cell populations, while it inhibited the LPS-induced production of the anti-inflammatory cytokine IL-10 (Table 1).

**Table 1.** Effect of prolonged 10 nM TSA exposure on pro- and anti-inflammatory cytokines release in LPS-stimulated microglia and astrocytes. Primary cultures of purified microglia and astrocytes were incubated with 10 ng/ml LPS in absence or simultaneous presence of 10 nM TSA for prolonged time exposure (24 hours). Then, TNF-α, IL-1β and IL - 10 releases were measured. The results are expressed as a percentage of LPS. Data are mean ± SEM of two or three independent experiments. Statistical analysis was performed with one-way ANOVA followed by Tukey’s test \**p*<0.05, \*\**p*<0.01 LPS + TSA 10 nM *vs* LPS.

|  | Astrocytes |  | Microglia |  |
| --- | --- | --- | --- | --- |
|  | LPS 10 ng/ml | LPS 10 ng/ml<br>+<br>TSA 10 nM | LPS 10 ng/ml | LPS 10 ng/ml<br>+<br>TSA 10 nM |
| **TNF-α** (% of LPS) | 100 ± 21.3 | 168 ± 12.1** | 100 ± 15.5 | 320 ± 4.2** |
| **IL-1β** (% of LPS) | 100 ± 10.2 | 149 ± 9.9* | 100 ± 16.2 | 259 ± 8.6* |
| **IL-10** (% of LPS) | 100 ± 18.8 | 39 ± 5.7* | 100 ± 0.9 | 76 ± 19.7 |

These results suggest 10 nM TSA modulates, over the LPS induction, the inflammatory response in both, astrocytes, and microglia.

### 3.2 Influence of the TSA and/or LPS pre-treatment schedule on cytokines release in mixed glial cells

Primary cultures of glia cells were also exposed to a) TSA (0.1 – 10 - 100 nM) for 1h and subsequently to 10 ng/ml LPS for 24h or b) LPS 1h before TSA (0.1 – 10-100 nM) for 24h (Fig 2 A). At the end of each exposure TNF-α was measured as representative cytokine (Fig 2 B).

**Figure 2.**
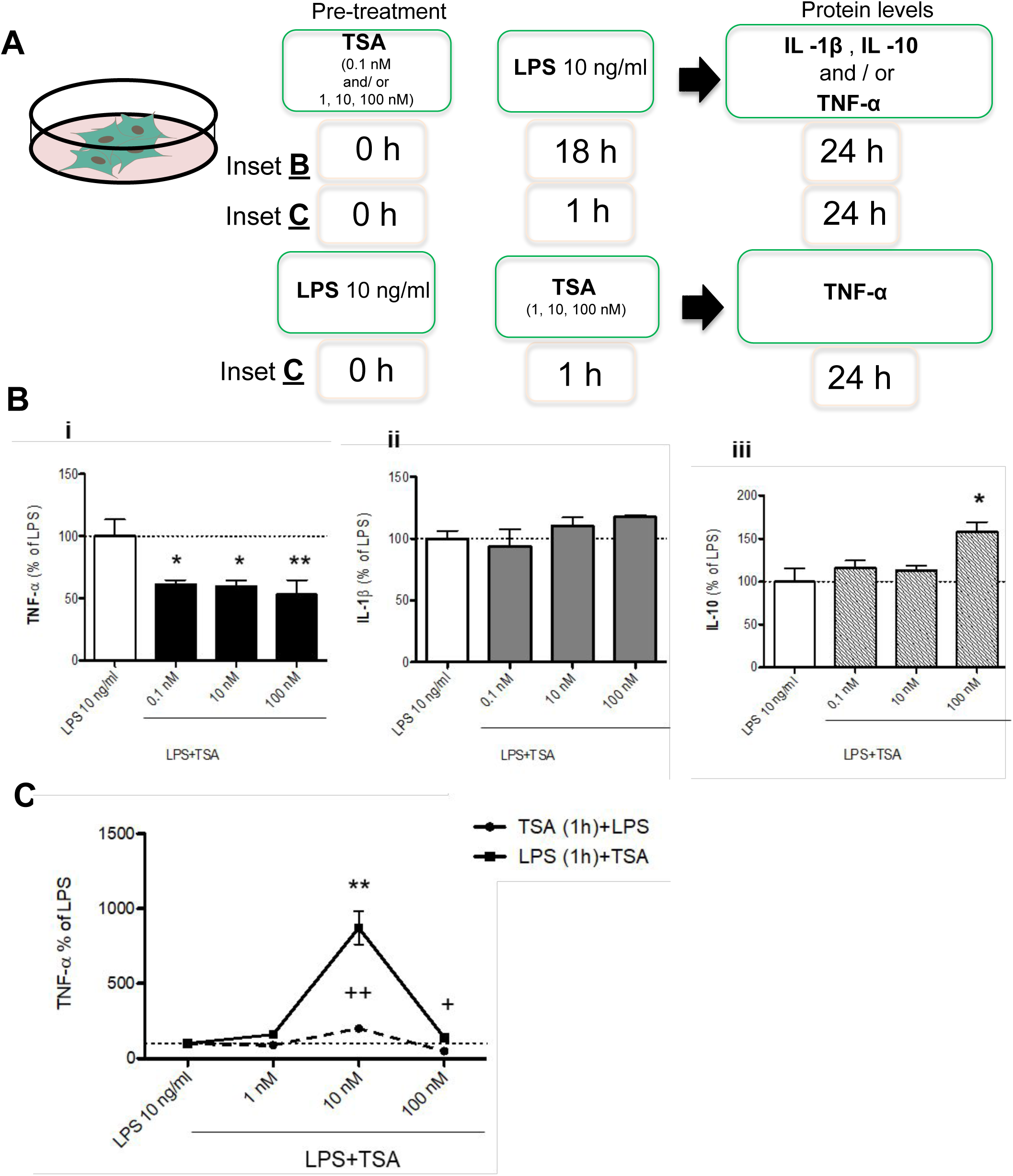
Influence of the TSA or LPS pre-treatment schedule on cytokines release in glia culture. **A) Experimental design.** Glia cells were pre-treated with different concentrations of TSA (0.1, 1, 10, 100 nM) and then 10 ng/ml LPS was added after 18 hours (TSA long-time exposure), (**inset B**), or after 1h (TSA short time exposure), (**inset C**). Glia cells were also pre-treated with 10 ng/ml LPS and then TSA (1, 10, 100 nM) was added after 1 hour (**inset C**). **B)** Protein levels of TNF-α **(i)**, IL - 1β **(ii)** and IL - 10 **(iii)** were measured in the culture medium after 24h by biological assay or ELISA for the inset B described above. The results are expressed as a percentage of LPS. **C)** TNF - α release after short pre-treatment (1h) with TSA (0.1, 10, 100 nM) or LPS (10ng/ml), (inset C described above). Data are mean ± SEM of two or three independent experiments in triplicate. Statistical analysis was performed with one-way ANOVA followed by Tukey’s test, \**p*<0.05, \*\**p*<0.01, LPS+TSA *vs* LPS; ++ *p*<0.01, TSA+LPS vs LPS.

Short pre-treatment at 10 nM TSA (a) significantly increased the production of TNF-α, while the highest dose, 100 nM TSA, significantly reduced the production of the selected cytokine (Fig 2 B). The increased TNF-a production induced by TSA 10nM is further enhanced with the pre-treated with LPS in mixed glia cells. No effect was observed for 1 and 100 nM TSA (Fig 2 B).

Primary cultures of glia cells were also exposed to TSA (0.1 – 10 - 100 nM) for 18h (Fig 2 A) followed by 10 ng/ml LPS treatment. TNF-α, IL-1β and IL-10 release were evaluated after 24h (Fig 2 C). By using this paradigm, the production of TNF-α at all tested concentrations was significantly reduced (Fig 2 C - inset i). On the other hand, IL-1β and IL-10 were not significantly affected, except for IL-10 at the highest dose of TSA, 100 nM (Fig 2 C - inset ii and iii respectively).

### 3.3 Dose-dependent effects of TSA on the expression of pro- and anti-inflammatory cytokines in LPS-co-stimulated primary alveolar macrophages

Primary cultures of alveolar macrophages were simultaneously treated with 10 ng/ml LPS in the absence or the presence of increasing concentrations of TSA (0.01, 0.1, 1, 10 nM) for 24 hours (Fig 3 A). The selected concentration of LPS did not affect macrophage viability (cell viability: control = 100% ± 2.2, LPS 10 ng/ml = 94.7% ± 12.1) as well as all the TSA tested concentrations, with or without LPS 10 ng/ml (cell viability referred to the highest concentration: control = 100% ± 3.6, TSA 10 nM = 81.50 ± 10.5, LPS 10 ng/ml + TSA 10 nM = 94.1% ± 1.20). Differently from what observed in glial cells, co-exposure of primary alveolar macrophages to increasing concentrations of TSA and LPS reduced both the production of pro-inflammatory (TNF-α, IL-1β) and anti-inflammatory (IL-10) cytokines compared to alveolar macrophages treated with LPS alone (Fig 3 B).

**Figure 3.**
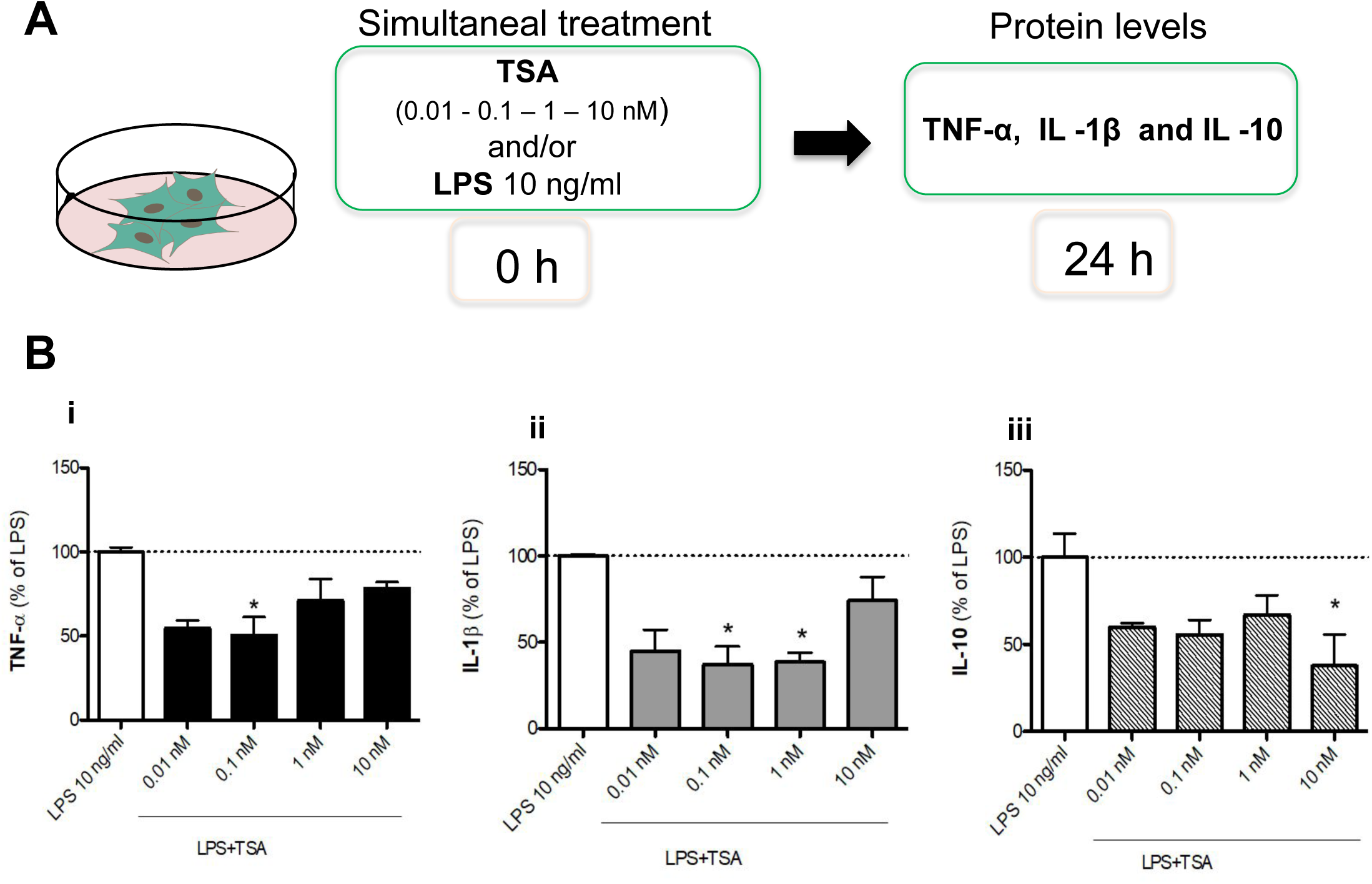
Effect of prolonged exposure to TSA on cytokines release in LPS-stimulated macrophages. **A)** Experimental design for prolonged TSA exposure. **B)** Primary cultures of alveolar macrophages treated with 10 ng/ml LPS in absence or simultaneous presence of different concentrations of TSA (0.01, 0.1, 1, 10 nM) for prolonged time exposure (24 hours). Protein levels of TNF - α **(i)**, IL - 1β **(ii)** and IL-10 **(iii**), released in the culture medium were measured by biological assay or ELISA. The results are expressed as a percentage of LPS. Data are mean ± SEM of two or three independent experiments in triplicate. Statistical analysis was performed with one-way ANOVA followed by Tukey’s test, \**p*<0.05, LPS+TSA *vs* LPS.

### 3.4 Dose-dependent effects of SAHA on the release and the expression of cytokines in LPS- stimulated glia cells

Primary cultures of mixed glia cells were treated simultaneously with 10 ng/ml LPS in the absence or the presence of increasing concentrations of SAHA (0.01, 0.05, 0.1, 0.5, 1, 2.5, 5 μM) for 24 hours and TNF-α release measured as previously described (Fig 4 A). All the tested concentrations of SAHA, with or without LPS, did not affect the glial cell viability (data not shown). Nanomolar concentrations of SAHA (50, 100 nM) significantly enhanced LPS-induced release of TNF-α, whereas micromolar concentrations of SAHA (2.5, 5 μM) significantly reduced it (Fig 4 B, inset i). Most prominent concentrations, 100 nM and 5 μM SAHA, were selected to co-expose glial cells with 10 ng/ml LPS for 24 hours and measure IL-1β and IL-10 release. No dualistic effect was observed as both nanomolar and micromolar concentrations of SAHA potentiated IL-1β, although to a lesser extent at 5 μM, and reduced IL-10 stimulated by LPS from glial cells. (Fig 4 B, inset ii and inset iii). A SAHA dose-response effect is evident for IL-10 reduction while SAHA 100 nM was most effective in potentiating IL-1 β (Fig 1, inset ii and inset iii).

**Figure 4.**
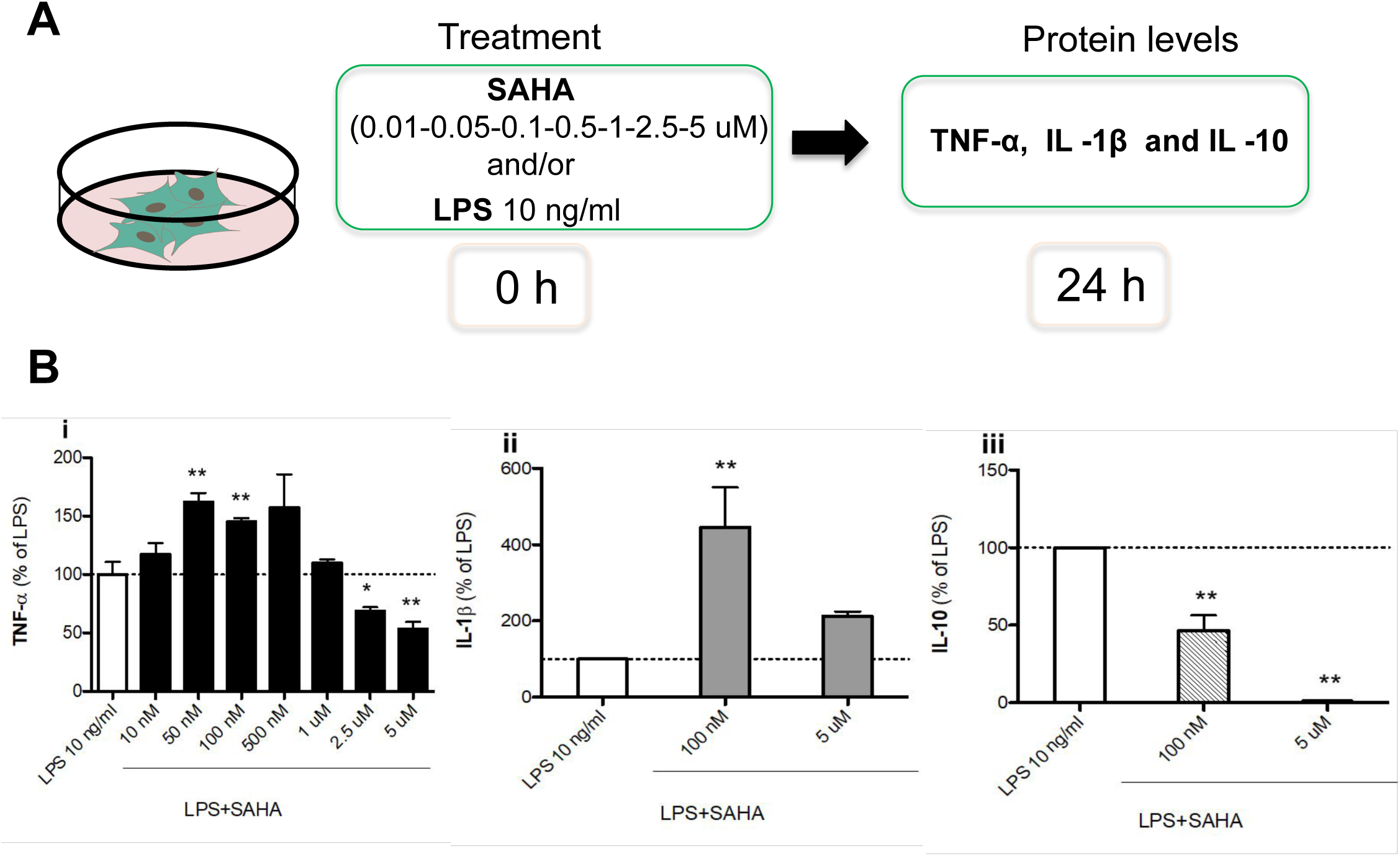
Effect of prolonged exposure to SAHA on cytokines release in LPS-stimulated glial cells. **A) Experimental design.** Glia cells were treated with 10 ng/ml LPS in absence or simultaneous presence of different concentrations of SAHA during long-time exposure (24 hours). **B)** TNF - α (SAHA: 50, 100, 500 nM, 1, 2.5, 5 mM) **(i)**, IL - 1 β (SAHA: 100 nM and 5 mM) **(ii)** and IL - 10 (SAHA: 100 nM and 5 mM) **(iii)** release was measured in the culture medium by biological assay or ELISA. The results are expressed as a percentage of LPS. Data are mean ± SEM of two or three independent experiments in triplicate. Statistical analysis was performed with one-way ANOVA followed by Tukey’s test, \**p*<0.05, \*\**p*<0.01 LPS+SAHA *vs* LPS.

### 3.5 Dose-dependent effects of Class I or II HDAC inhibitors on the expression of pro- and anti-inflammatory cytokines in LPS-co-stimulated mixed glia cells

Using the previous paradigm condition, cells were simultaneously exposed to 10 ng/ml LPS in the absence or the presence of increasing MS-275 (class I HDAC inhibitor) or MC1568 (class II HDAC inhibitor) concentrations for 24 hours (Fig 5 A). Then, TNF-α release was measured in the culture medium. MS-275 significantly reduced TNF-α released by LPS in a dose-dependent manner (Fig 5 B, inset i), while MC1568 enhanced it (Fig 5 C, inset i). Primary cultures of glia were treated with 10 ng/ml LPS in the absence or the presence of 100 nM MS-275 or 5 μM MC1568, as representative concentrations, the production of IL-1 β and IL10 was measured after 24 hours from the exposure. As reported in figure 5 B and 5 C (inset ii and iii), 100 nM MS-275 did not affect the production of the selected inflammatory molecules. Differently, 5 μM MC1568 potentiated the LPS-induced release of IL-1β and significantly reduced the production of IL-10 in LPS-stimulated glia cells (Fig 5 B and C).

**Figure 5.**
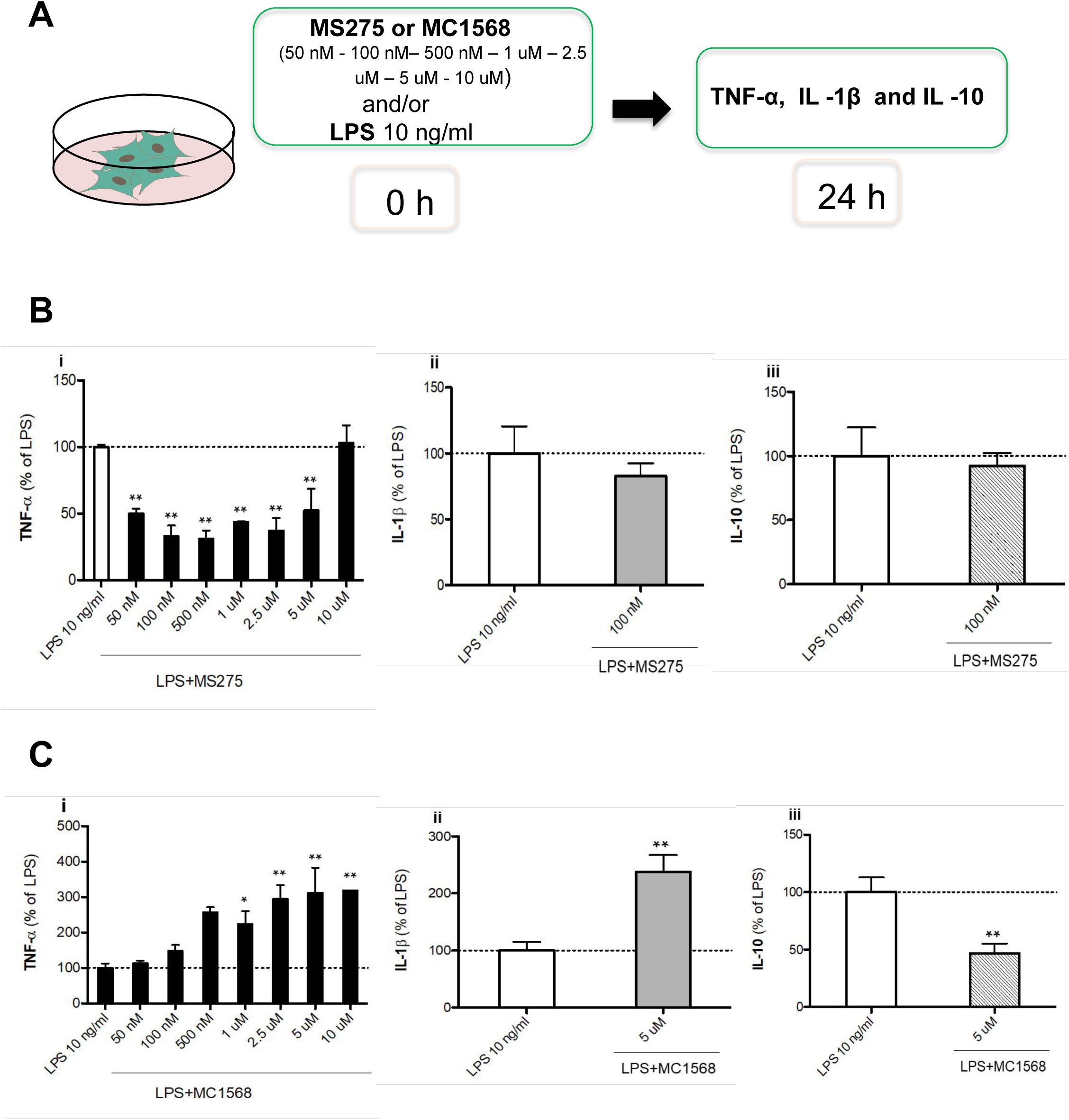
Effect of selective class I HDACs (MS275) and class II HDACs (MC1568) on cytokines release in LPS-stimulated glial cells. Glia cells treated with 10 ng/ml LPS in absence or simultaneous presence of MS275 **(B)** or MC1568 **(C)** during long-time exposure (24 hours). TNF-α (MS275 and MC1568: 50, 100, 500 nM, 1, 2.5, 5, 10 mM) **(i)**, IL-1β (MS275: 100 nM; MC1568: 5 mM) **(ii)** and IL - 10 (MS275: 100 nM; MS275: 5 mM) **(iii)** release was measured in the culture medium by biological assay or ELISA, inset **A)** and **B)**. The results are expressed as a percentage of LPS. Data are mean ± SEM of two or three independent experiments in triplicate. Statistical analysis was performed with one-way ANOVA followed by Tukey’s test, \*\**p*<0.01 LPS+ MS275 or MC1568 *vs* LPS.

### 3.6 DNA microarray profiles of genes differentially regulated by 100nM and 5μM SAHA

#### 3.6.1 Gene expression modulation after co-exposure of glia cells to LPS and 100nM or 5μM SAHA

To better understand the dualistic effects of SAHA, microarray analysis of gene expression profile of glia cells was performed after co-exposure of cells to 10 ng/ml LPS and 100 nm, 5 μM SAHA, or vehicle (LPS) for 24 hours.

Transcription profile was compared between LPS-induced cells treated with 100 nM SAHA (LPS_SAHA100 – LPS) and LPS-induced cells treated with 5 μM SAHA (LPS_SAHA5 – LPS), (Fig 6 A and B). A total of 97 differentially expressed genes (DEGs) were differentially expressed by LPS_SAHA100. 17 genes were significantly upregulated, while 80 genes were significantly downregulated. Instead, LPS_SAHA5 showed a much stronger impact on transcription, with 1628 DEGs, 744 upregulated and 884 downregulated, respectively (Fig 6 A).

**Figure 6.**
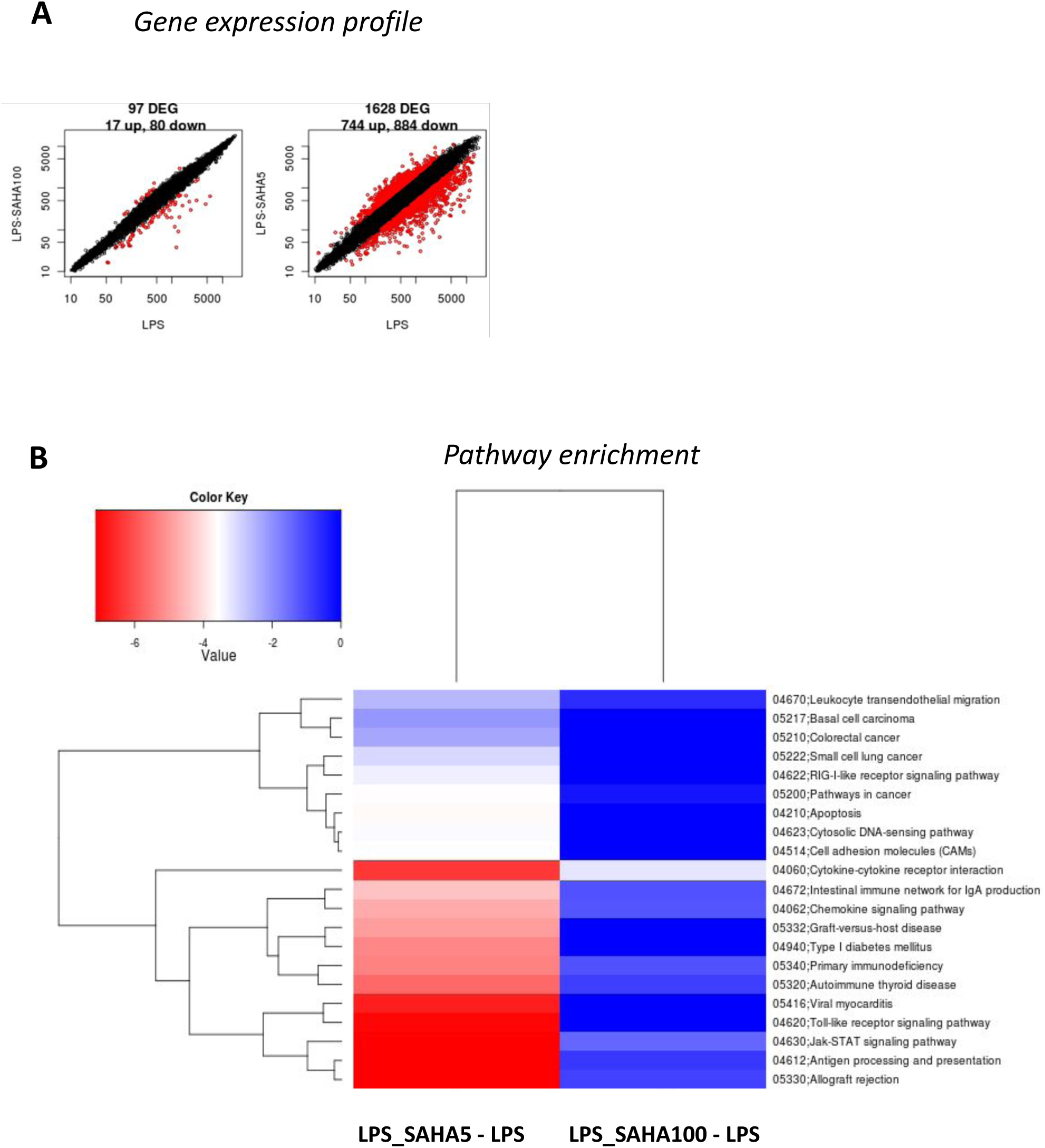
A) Scatter plots of log2 intensity values of DEGS. Total number of DEGs (red dots) identified for glia cells treated with SAHA 100 nM or SAHA 5 uM compared to LPS (10 ng/ml) condition. DEGs identified with the Limma algorithm with a threshold value of 0.01. On the y-axis are reported the means of the replicates of the treatment conditions and on the x-axis the mean across the replicates of the reference condition (LPS 10 ng/ml). **B) Pathway enrichment analysis.** KEGG functional summary reporting the enrichment p-values less than 0.05 (log10 of p-value) for the different annotation terms LPS_SAHA100 - LPS and LPS_SAHA5 - LPS. The color scheme is based on a simple scale (reported above) painting pathways (blu down – and red up-regulated) by automatic analysis using Basic Analysis.

Notably, only few DEGs were shared between the two concentrations of SAHA (Table S1). Among them, genes involved in immune response modulation (i.e., *Fap, Gcg, Ccl22, Slamf1, Cxcl13*) were strongly repressed, while signal transduction pathways were highly increased (i.e., *Plekhh*). Genes negatively regulated uniquely for LPS_SAHA100 – LPS comparison were also reported, as *Gpr183, Rgs2, Rgs1* (gene involved in regulate G-protein signalling), as well as for LPS_SAHA5 – LPS, *Clec4e, Clec4e, Has2, Ccr5* (genes involved in signalling receptor activity for inflammatory and immune response). On the other hand, genes positively regulated exclusively for LPS_SAHA100 – LPS were emerging, i.e. *LOC302473, Zfp57, Rreb1, Zbtb7b* (genes involved in transcription regulation for the immune response), as well as genes for LPS_SAHA5 – LPS comparison, *Gdf15, Plag1* (genes involved in cytokine and transcription factor activity). Finally, only Mmp9 showed opposite behaviour in response to low and high SAHA concentration (up in LPS_SAHA100 and down with LPS_SAHA5, respectively).

Gene expression profiles were then compared through analysis of canonical pathways enrichment, KEGG, as already described in Materials and Methods. Enrichments significantly modulated, according to DEG functional annotation analysis, are reported in Fig 6 B.

Most of these pathways were modulated according to the dose used and mainly by high SAHA concentration (LPS_SAHA5). We then examined only pathways significant altered by both comparisons (“Cytokine-cytokine receptor interaction” and “JAK-STAT signalling pathway”), and we looked at specific DEGs in these pathways (Table 2), to identify possible opposite transcript regulations due to the dose. We found few DEGs commonly altered and genes where the expression was only regulated in the comparison LPS_SAHA5 - LPS, Moreover, a significant down-regulation of IL-10 gene is shown for both doses and in all the pathways considered, confirming our previous molecular results in glia cells.

**Table 2.**
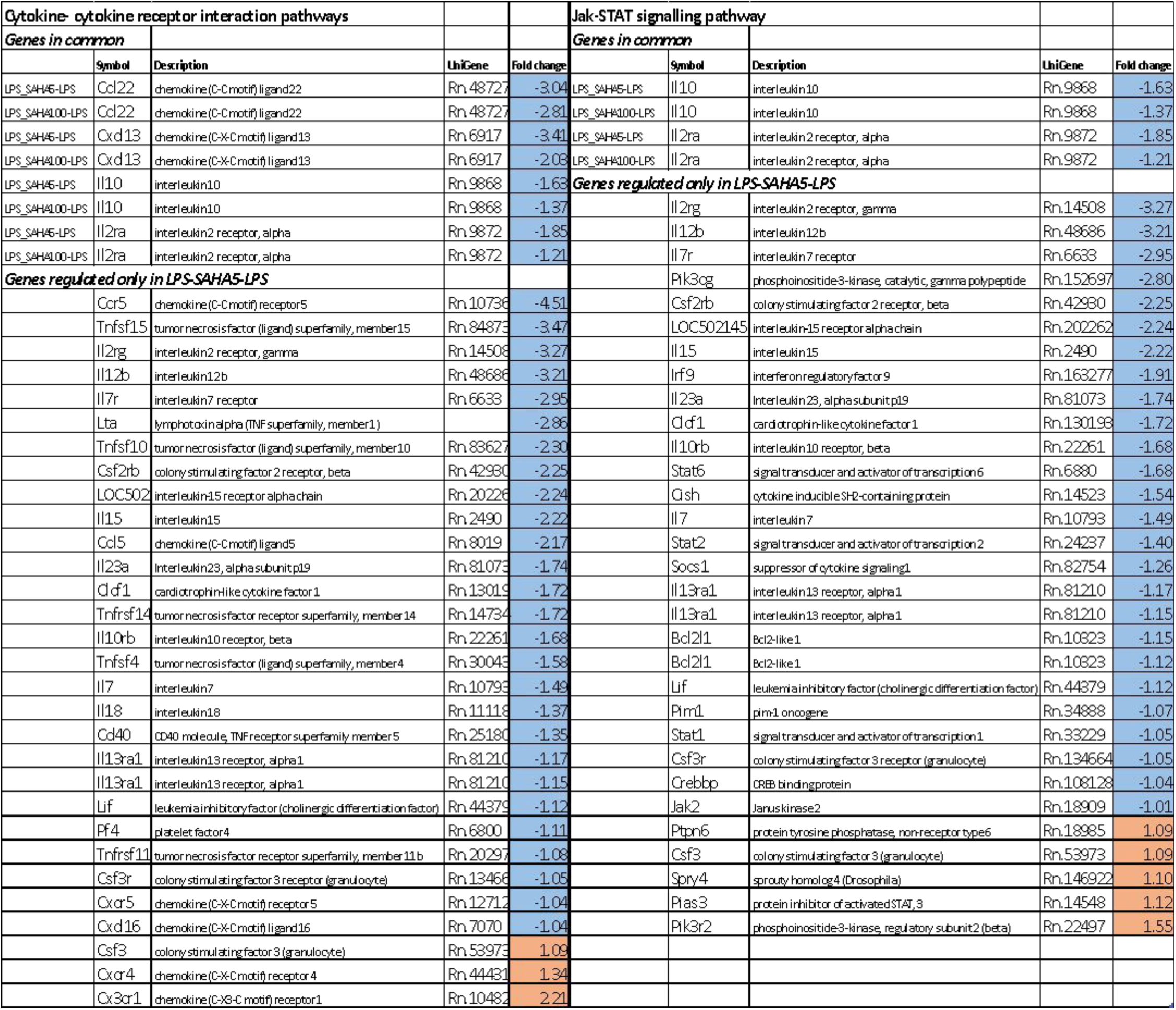
DEGs members were detected as significantly expressed for the condition LPS_SAHA5 - LPS and LPS_SAHA100 – LPS for the pathways “Cytokine - cytokine receptor interaction” and “JAK / STAT signalling pathway”. Color schema follows the statistic of differentially expressed (blue stands for down – and red for up-regulated genes, NA-not detected).

To gain insight into the pathways and networks affected by treatment conditions, analysis was also performed with gene ontology (GO) enrichment (Fig S1). Pathways related to chemokine activity, cytokine receptor activity and immune system were differently regulated between treatments and mainly upregulated by the higher dose of SAHA.

#### 3.6.2 Gene expression modulation on control condition of mixed glia cells

To see the impact of the treatment on glia cells without a prior LPS inflammatory stimulus, transcription profiles were analysed for SAHA100 - CTRL and SAHA5 - CTRL groups (Fig S2 and S3).

For the comparison SAHA100 - CTRL, a total of 18 DEGS were differentially expressed, two of which were upregulated while 16 were repressed. A stronger effect on transcription was observed with high SAHA concentration (SAHA5). Specifically, 849 genes were differentially modulated in response to 5 uM SAHA (339 upregulated and 510 downregulated, respectively); (Fig S2 A). Only a few genes were shared between the two conditions (Table S2). Among them, genes involved in inflammatory response such as Neuropeptide Y (*Npy*), Interferon-induced protein 1 (*Ifit1*), Ermin (*Ermn*) and GNAS (*Gnas*) showed dose-dependent downregulation in response to SAHA. Consistently, KEGG pathways, biological process (BP), molecular function (MF), and cellular components (CC) analysis showed significant enrichment in inflammatory processes and immune response (Fig S2 and S3, respectively).

#### 3.6.3 Gene expression modulation for LPS-induced and control treatment in mixed glia cells

To see differences due to the treatments on control cells and LPS-induced cells, DEGS were analysed, and a Venn diagram was generated (Fig S4). Venn diagram reports two sets of comparisons: a) *SAHA 100 - CTRL* and *SAHA 100 - LPS* and b*) SAHA 5 - CTRL and SAHA 5 - LPS.* The intersection part gives a direct view on how many genes are involved in possibly related functions. For the condition a) treatment with SAHA 100, 14 genes were found to be modulated when SAHA was added to control cells, 93 genes modified their expression when added in presence of LPS. Only 4 genes were modified by SAHA both in the presence and the absence of LPS. For the condition b) treatment with SAHA 5, 335 genes were significantly modulated in the absence of LPS, 1114 genes in the presence of LPS, and 514 in both conditions (Fig S4).

DEGs significantly modulated in the intersection between comparisons are reported in Table S3. Among these, for the condition a), down-regulation of *Npy* (neuropeptide Y) was reported (the other three genes were not identified, NA). For the condition b), genes, with lower fold change value, emerged, such as *Npy*, *Nav3* (neuron navigator 3), *Fst* (follistatin) while *Napepld* (N-acyl phosphatidylethanolamine phospholipase D), *Plag1* (Pleiomorphic adenoma gene 1) presented the highest positive fold change value. All these genes are involved in regulating neuroinflammation via modulation of cellular homeostasis and signal transduction. Interestingly, *Npy* was reduced with both SAHA concentrations. This finding could be related to the pro-inflammatory action exerted by *Npy*. Indeed, key signaling pathways in immune cells activated by neuropeptides include nuclear factor-κB (NF-κB), cyclooxygenase-2 (COX-2), or mitogen-activated protein kinase (MAPK) that further propagate inflammatory signals [38].

## DISCUSSION

Previous evidence reported that the inhibition of HDAC activity regulates inflammatory response in activated microglia. However, contrasting results emerged for mouse and rat microglia [23] [27][20][24] [43] and according to the type of cells used, i.e. primary mouse dendritic cells [26]. As such, the mechanism by which HDAC inhibition is beneficial or harmful in glia is still far from being completely elucidated.

This study demonstrated different effects of HDACi on inflammatory response according to:

a. the concentration of the HDACi used.
b. the paradigms of treatment used (simultaneous or prior treatment with pro-inflammatory stimulus, acute or chronic exposure);
c. the cell type (mixed glia cells, purified microglia and astrocytes, alveolar macrophages
d. the inhibition spectrum achieved: broad-spectrum inhibitors (TSA and SAHA) vs. class-selective inhibitors (MS275 per class I HDACs, and MC1568 for class II HDACs).

The impact of the different experimental conditions used on the LPS response has been evaluated through the expression and release of a set of pro-inflammatory (TNF-α, IL-1β) and anti-inflammatory (IL-10) cytokines. As reported from our results, TNF-α is the only cytokines whose release is modulated by all the HDACs inhibitors considered.

We also showed that these effects involved regulation of transcripts of different homeostatic pathways and that this regulation is highly dependent on the dose of HDACi used, with 5 µM SAHA being the most active dose in this mechanism.

Here, we demonstrated that the concomitant exposure to LPS and TSA, strongly potentiates the inflammatory response in primary culture of mixed glial cells, being more evident at the concentration of 10 nM. However, TSA at an increased dose of 100 nM significantly reverts the previous response, with a strong enhancement of IL-10 release. This effect is not shared with SAHA and could be specific for TSA, as it was already described the TSA ability to elevate Il10 gene expression levels by increasing histone H3 acetylation recruitment [44]]. Indeed, at similar concentration (100 nM) TSA increased IL-10 release while SAHA decreased it. In similar conditions, our results also reported a more efficient effect of TSA in producing cytokines releasing at the dose of 1nM compared to SAHA, whose effect is evident at 50nM. Moreover, short time exposure (1h) to 10 nM TSA increases LPS-induced expression of the immediately pro-inflammatory transcripts, *TNF-α* and *IL-1β* and reduces the expression of *IL-10*, indicating the ability of TSA to regulate earlier neuroinflammatory response at transcriptional level.

Notably, long-time exposure of glia cells to TSA before the inflammatory stimulus, reduces TNF-α, increases IL-10, and does not affect IL-1β. On the contrary, short pre-treatment (1 hour) with 10nM TSA potentiates inflammatory response in terms of TNF-α release. These last data suggest that HDACi leads to the activation of glial cells independently of LPS pre-treatment. A similar effect in TNF-α, release, was also observed with a short pre-treatment with LPS, although less conspicuous. In this view, evidence reported the ability of TSA to enhance the inflammatory effects of LPS when added simultaneously [30]. However, pre-treatment with 15nM of TSA showed an anti-inflammatory effect in microglia cells [20], (data not shown). Indeed, TSA differently modulates LPS-induced inflammatory response of glial cells, depending on treatment protocol [45]. These observations reveal that paradigms of exposure and the dose used can influence the ability of HDACi to modulate the inflammatory response, helping to complement and clarify the wide discrepancy in literature related to the effects of HDAC inhibition on neuroinflammatory response.

We also wanted to test the ability of TSA to inhibit early inflammation response on different cell populations. The pro-inflammatory effect of TSA on LPS-induced neuroinflammation occurs also in primary cultures of purified astrocytes and microglia at the dose of TSA 10 nM upon 24h exposure. The level of cytokine induction in astrocytes in response to LPS was lower than in microglia, supporting the notion that the innate immune response is mediated primarily by microglia [46][47]. Differently from what is observed in glia cells, in the primary culture of macrophages, simultaneous treatment with LPS and TSA leads to an anti-inflammatory response, reducing the production of both pro- and anti-inflammatory cytokines induced by LPS. Some reports highlighted that broad-spectrum HDACi, such as SAHA, diminished the production of key pro-inflammatory cytokines from LPS-stimulated peripheral blood mononuclear cells [48]. Similar anti-inflammatory effects were reported in subsequent studies [25][49] where TSA inhibited the effects on the nitric-oxide (NO) production in bone-marrow-derived macrophages [50]. Our results are in line with previous evidence [51] [52][53] suggesting that the effect of HDAC inhibition on inflammatory response may depend on the cell type. Another explanation for these opposite effects of HDACi on inflammatory response could also be attributed to the differential role of individual HDAC isoforms in cellular functions and to the non-specificity of the HDACi used, as previously reported [54].

In this regard, we also tested selective inhibitors of class I and class II using the same treatment conditions (24h). Our results showed that simultaneous treatment of glia cells with LPS and the selective class I HDACi MS275 reduces TNF-α release. On the contrary, the class II HDAC-selective inhibitor MC1568 dose-dependently increases TNF–α production. Several studies showed that class I HDACi MS275 reduces neuroinflammation in a mouse model of Alzheimer’s disease [55][56][57] and suppresses cytokine expression in murine microglia activated with LPS [43]. Although several pieces of evidence reported the MC1568 ability to modulate inflammation according to the cell types [58][59][60], to the best of our knowledge, other than our data, there is no evidence of MC1568’s ability to modulate the glial inflammatory response.

For the foregoing, our results highlight the importance of the dose on the dualistic effects of these drugs in inflammatory response and the ability of all the pan inhibitors tested to have an important effect in the modulation of TNF-α release.

Same experimental conditions (prolonged exposure, 24h) were used to test the ability of the broad spectrum HDACi SAHA in modulating glia inflammatory processes triggered by LPS. The concomitant treatment with 10 ng/ml LPS and SAHA strongly potentiates the inflammatory response in primary culture of mixed glia cells at the concentration 100 nM. However, at 5 µM SAHA significantly reverts the previous response on cytokine release. Several studies already reported the anti-inflammatory properties of SAHA in mouse glia cells [61], where LPS-induced TNF-α and IL1 protein expression was inhibited by 1 µM SAHA [47].

Although this neuroprotective action, the underlying mechanisms of SAHA in the inflammatory process are still uncertain and may depend on the SAHA’ s ability to modulate numerous host cell transcripts.

Within this frame, our transcriptomic analysis identified patterns of gene expression across different experimental conditions with SAHA treatments. Different dose of SAHA, 5uM and 100 nM, predominately results in the transcriptional dysregulation of JAK signalling, potentially by regulation of *JAK2* gene, as well as altered activation of cytokines cascade signalling mainly through expression of *IL-10* gene, respectively. Indeed, only two signalling pathways were differentially modulated by both doses of SAHA: “JAK/STAT pathway” and “Cytokines-cytokines receptor interaction pathways”. In accordance with our molecular results, our transcriptome analysis showed that both doses of SAHA decrease the expression of *IL-10* interleukin. Although we did not find any direct modulation in the expression of *TNF-α* and *IL-1β,* we observed alteration of key transcripts involved in *TNF-α*, *IL-1β* and IL-10 downstream pathway activation. Our transcriptome results may suggest an indirect modulation of the maturation of cytokines through the regulation of the expression of key factors involved in the downstream pathways. Specifically, for the LPS_SAHA5 and LPS comparison, several members of the tumour necrosis factor receptor superfamily were found to be negatively regulated while the expression of *Lta,* and *Casp-8* while *Traf4* and *Fadd* were upregulated. These genes are important in inflammation and cell death pathways. Indeed, caspase-8 can form a complex with the adaptor FADD, along with additional constituents, inducing cytokine production in response to apoptotic stimuli [39]. Moreover, it has been shown that TNF-α promotes apoptosis in astrocytes through the RIP1/FADD/Caspase-8 axis [40].

Another example is CASP1, a protein responsible for converting pro-IL-1β to the active IL-1β [62], and its transcript appeared downregulated in our condition with the dose of 5µM SAHA, emphasising on the anti-inflammatory effects at SAHA 5µM. In addition, for the LPS_SAHA5 - LPS comparison, we also found a decrease in the expression of interleukin 10 receptor beta, *Il10rb*, along with concomitant upregulation of the protein inhibitor of activated *Stat3, Pias3*. These genes are in the downstream pathway of IL-10 being STAT3 protein activated by the IL-10 family of cytokines, and inhibited by PIAS3, thus mediating the anti-inflammatory response [42].

Contrary to 5µM SAHA, with 100 nM SAHA we observed only a downregulation of *IL10* and *IL2Ra* (receptor antagonist of IL-1β). Evidence reported that IL-10 plays an important role as a negative regulator for cytokine production in microglia culture [63], by suppressing STAT3-caspase 1 dependent IL-1β maturation [64] and exerting, thus, the anti-inflammatory action [65]. In this view and based on our transcriptomic results, we hypothesized that dysregulated IL10-STAT3-IL-1β signalling feedback could be responsible for the increased inflammatory cytokines with 100 nM SAHA. Based on transcriptomic analysis, other mechanisms are likely to partake of the anti-inflammatory action of the high dose of SAHA, including JAK-STAT signalling.

The “JAK/STAT pathways” represents a well characterized cell signalling system, playing an important role in disease states, particularly neuroinflammatory diseases [66].

Different JAKs and STATs proteins have been identified, and each cytokine exerts its function through a specific combination of JAK and STAT members [67] [68].

Here, we demonstrated a possible anti-inflammatory effect of 5 µM SAHA via modulation of key genes of this pathway. Indeed, downregulation of several *STATs (1, 2, 6), JAK2*, interleukins, interferons, *BCL2l1* and upregulation of genes generally involved in negative feedback, such as *PIAS, Sprouty, and P13K* [69], were observed.

It is worth mentioning that previous studies have shown the JAK2-STAT3 signalling is involved in microglia activation [70]. Moreover, STAT1-dependent gene activation is regulated by HDACs [71][72], meaning that the inhibition of HDAC activity prevents IFN γ–induced JAK1 and STAT1-dependent gene activation [73], playing a central role in the regulation of immune responses [69]. In this view, evidence supports the anti-inflammatory effects of SAHA through attenuation of the JAK/STAT pathway in myeloproliferative neoplasms [74] and cognitive dysfunctions [75].

In conclusion, our study revealed a variable response to HDACi depending on cell type, HDACi dosage, protocol and HDACi specificity used. Indeed, different concentrations of wide spectrum HDACi (e.g., TSA and SAHA) impact in a dose dependent-manner inflammatory response in glia cells through specific cellular processes at transcriptional and protein level. Our approach identified broad gene networks and functional pathways affected by SAHA; however, the biological significance of the individual genes identified should be validated by more focused and functional studies. Our global gene expression analysis under different conditions unveiled genetic pathways and provides the opportunity to highlight potential molecular events, underlying the role of clinically used HDACi in the prevention of neuroinflammation.

## SUPPLEMENTARY FIGURES

**Figure S1. GO functional summary** for the comparison LPS_SAHA100 or 5 - LPS. **A)** Biological Process (BP) **B)** Molecular Function (MF) **C)** Cellular Components (CC), reporting the enrichment p-values less than 0.05 (log10 of p-value).

**Figure S2. A) Scatter plots of log2 intensity values of DEGS.** Total number of DEGs (red dots) identified for glia cells treated with 100 nM or 5 uM SAHA compared to control (CTRL) condition. DEGs identified with the Limma algorithm with a threshold pvalue of 0.01. On the y-axis are reported the means of the replicates of the treatment conditions and on the x-axis the mean across the replicates of the reference condition (CTRL). **B) Pathway enrichment analysis.** KEGG functional summary reporting the enrichment p-values less than 0.05 (log10 of p-value) for the different annotation terms and their similarity for the comparisons SAHA100 - CTRL and SAHA5 - CTRL. The color scheme is based on a simple scale painting pathways (blu down – and red up-regulated) by automatic analysis using Basic Analysis.

**Figure S3. GO functional summary** for the comparison 100 nM or 5 uM SAHA - CTRL. **A)** Biological Process (BP) **B)** Molecular Function (MF) **C)** Cellular Components (CC), reporting the enrichment p-values less than 0.05 (log10 of p-value).

**Figure S4.** Treatment effects in gene expression modification for control cells and LPS-induced cells. Venn diagram with two sets **A)** *SAHA 100 - CTRL and SAHA 100 - LPS*, **B)** *SAHA 5 - CTRL and SAHA 5* - *LPS* and their intersections.

## SUPPLEMENTARY TABLES

**Table S1.** DEGs members were detected as significantly expressed for the condition LPS_SAHA100 - LPS and LPS_SAHA5 - LPS. Color schema follows the statistic of differentially expressed (blue stands for down – and red for up-regulated genes, NA-not detected).

**Table S2**. DEGs members were detected as significantly expressed for the condition SAHA100 – CTRL and SAHA5 - CTRL. Color schema follows the statistic of differentially expressed (blue stands for down – and red for up-regulated genes, NA-not detected).

**Table S3**. DEGs members are detected as significantly expressed in the intersection between SAHA 5 - CTRL and SAHA5 - LPS or SAHA 100 - CTRL and SAHA100 - LPS, Venn Diagram. Color schema follows the statistic of differentially expressed (blue stands for down – and red for up-regulated genes, NA-not detected).

## Notes

### Competing Interest Statement

The authors have declared no competing interest.

