## Supplementary figures and tables for "DUALISTIC SAHA DOSE-DEPENDENT EFFECTS ON GLIAL-INFLAMMATORY RESPONSE": Supplementary Figures 2 BV (4).pptm

### Slide 1
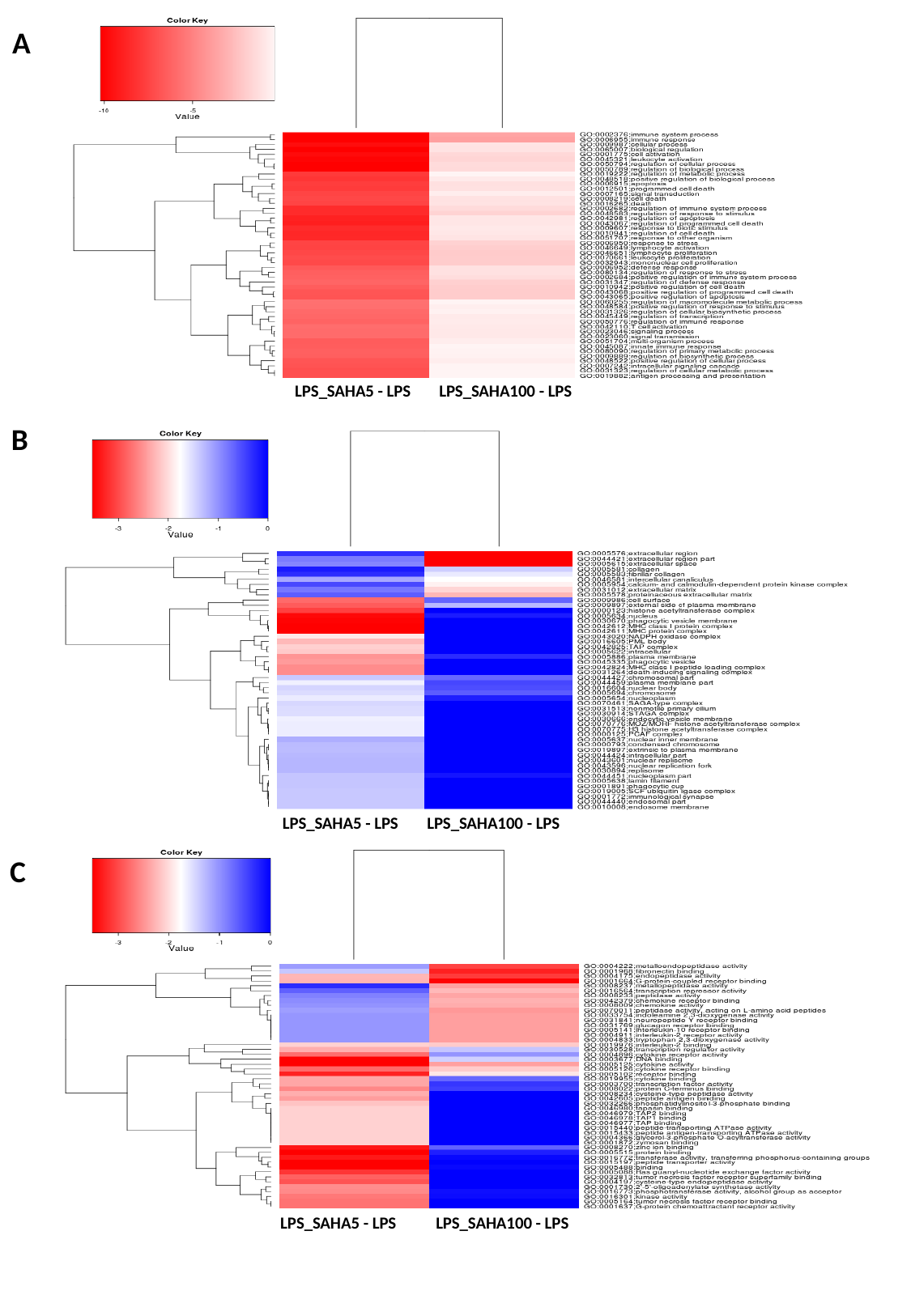

A
LPS_SAHA5 - LPS
LPS_SAHA100 - LPS
B
LPS_SAHA5 - LPS
LPS_SAHA100 - LPS
C
LPS_SAHA5 - LPS
LPS_SAHA100 - LPS

### Slide 2
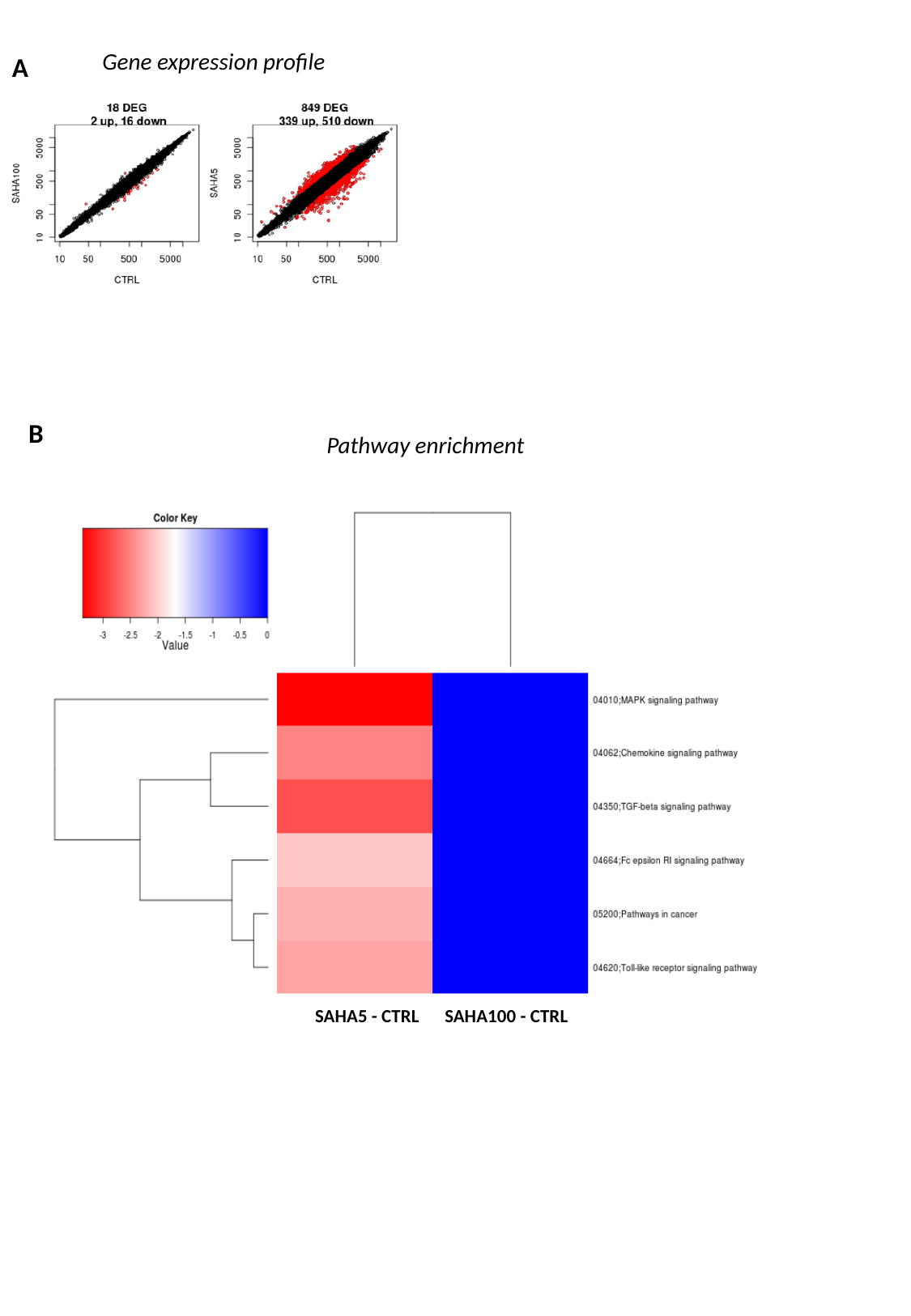

Gene expression profile
A
B
Pathway enrichment
SAHA5 - CTRL
SAHA100 - CTRL

### Slide 3
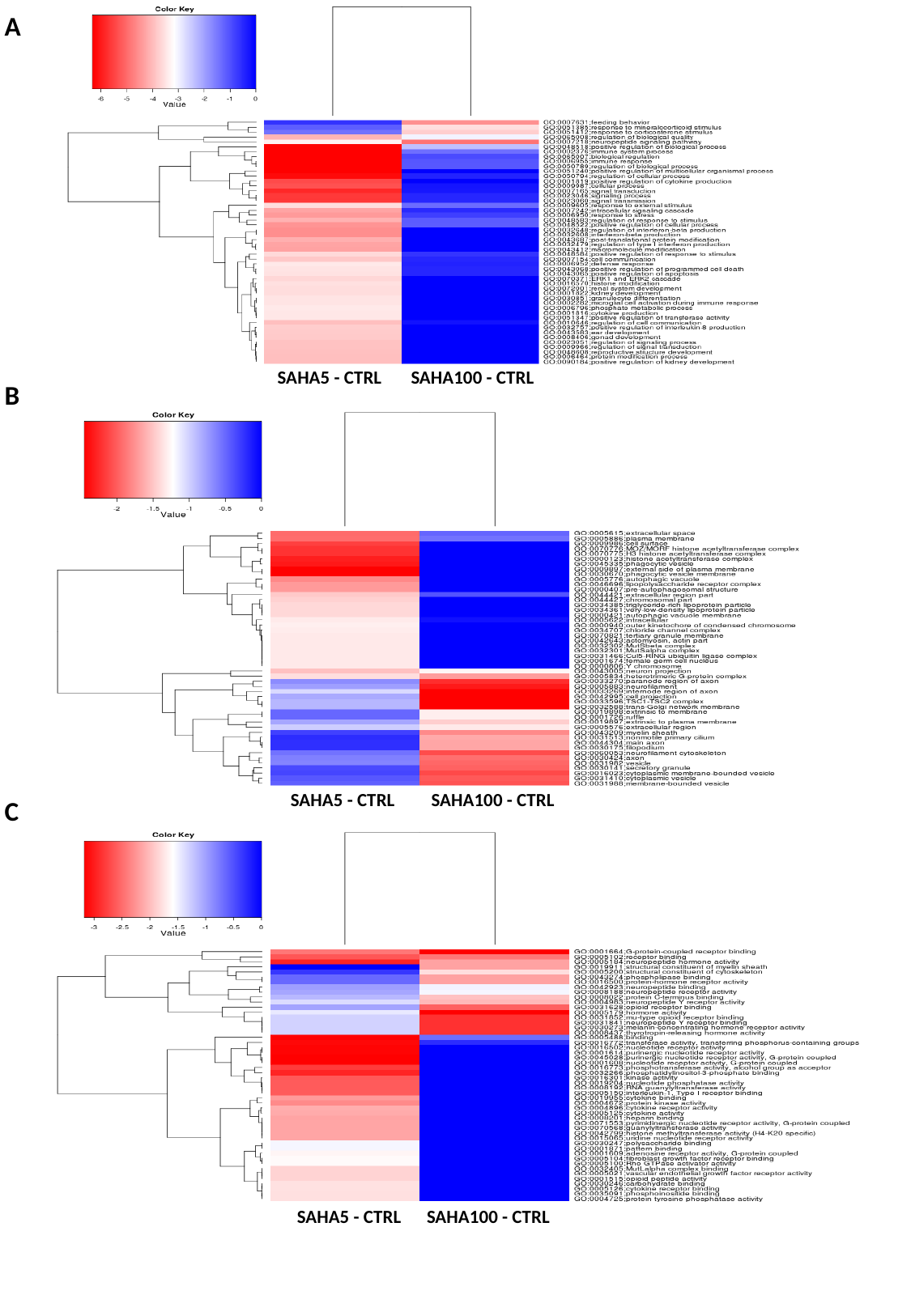

A
SAHA5 - CTRL
SAHA100 - CTRL
B
SAHA5 - CTRL
SAHA100 - CTRL
C
SAHA5 - CTRL
SAHA100 - CTRL

### Slide 4
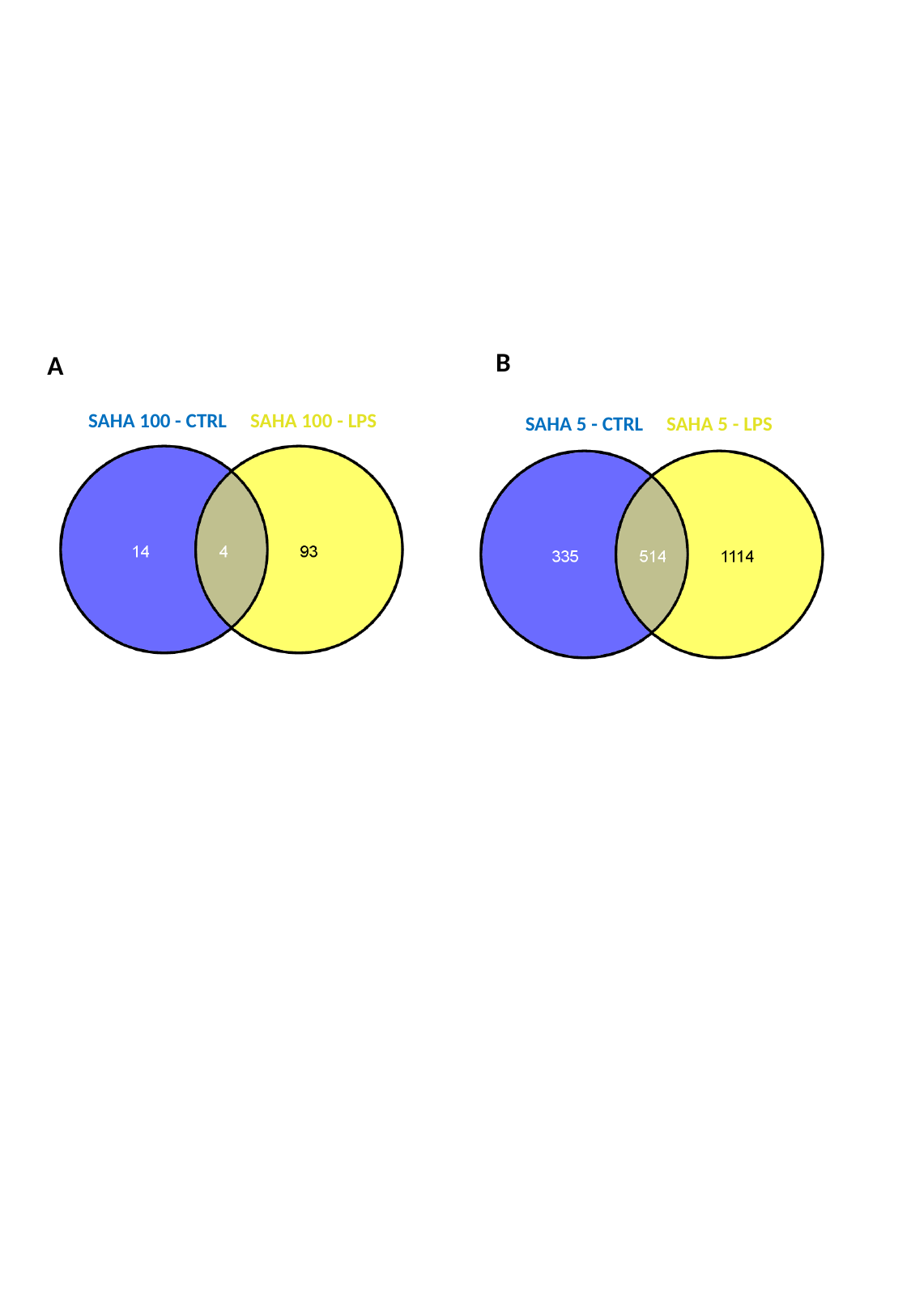

B
A
SAHA 100 - CTRL SAHA 100 - LPS
SAHA 5 - CTRL SAHA 5 - LPS
